# Adaptive Cancer Suppression among Tissues through Reduced Stem Cell Mutation Rates

**DOI:** 10.64898/2026.04.13.718347

**Authors:** Jack da Silva

## Abstract

Since most cancers are initiated by driver mutations arising in somatic cells, the risk of cancer should be explained by the probability of driver mutations as a function of the product of the number of stem cells and the stem cell division rate for a tissue. However, such models of driver-mutation initiated carcinogenesis fail to adequately predict cancer risk. It has been suggested that the missing component is greater adaptive cancer suppression in large tissues with high rates of stem cell division. This 8 may manifest as either a greater number of driver mutations required to initiate cancer or a lower stem cell mutation rate. These hypotheses are tested here using nonlinear regression to fit models that incorporate variation in the number of driver mutations and the stem cell mutation rate to data on the tissue-specific cancer risk, number of stem cells, and stem cell division rate. The greatest concordance between predicted and observed cancer risks is obtained by single driver mutations across tissues and stem cell mutation rates that decline with increasing lifetime numbers of stem cell divisions. This provides evidence of adaptive cancer suppression among tissues.

## Introduction

Most cancers are initiated by driver mutations that inactivate tumour-suppressor genes or activate oncogenes [1-3]. Some of these mutations may be inherited, but germline mutations for cancer are overwhelmingly recessive [4], likely as a result of strong selection against rare dominant mutations of large effect. Therefore, driver mutations often arise in somatic cells [5]. In humans, it is estimated that 66% of cancers are due to mutations arising during stem-cell division, 29% are due to somatic mutations caused by environmental factors, and 5% are inherited [6, 7]. This effect of stem cell division was estimated from the correlation between the total number of stem cell divisions over an individual’s lifetime for a given tissue and the lifetime risk of cancer in that tissue across 31 tissues/cancers. The main conclusion was that the probability of driver mutations arising increases with the lifetime number stem cell divisions. Thus, larger tissues, tissues with more rapidly dividing stem cells, larger bodies, and longer-lived individuals should be more susceptible to cancer. This is consistent with the observation that cancer risk increases with height in humans [8-13] and with weight and lifespan in dog breeds [14-19].

However, the lifetime number of stem cell divisions provides an incomplete explanation of the risk of cancer for a tissue due to mutations arising from stem cell division. Simple models of carcinogenesis that assume a constant number of driver mutations are required to initiate cancer across all tissues predict the risk of cancer should increase linearly with the lifetime number of stem cell divisions with a slope of one on log scales [20]. However, the risk of cancer actually increases with a slope significantly less than one on log scales. This may be explained if the number of driver mutations required varies among tissues due to varying levels of adaptive cancer suppression [21-23]. In other words, tissues that are more prone to mutation because of high numbers of stem cell divisions may evolve increased cancer suppression in the form of higher numbers of driver mutations required to initiate cancer. Alternatively, greater cancer suppression may be attained through increased inhibition and repair of DNA damage [24, 25], reducing mutation rates. Thus, variation in the risk of cancer among tissues may be further explained by a greater number of required driver mutations or a lower mutation rate in tissues with higher lifetime numbers of stem cell divisions.

Here, models of driver mutation-initiated cancer are fitted to the Tomasetti CVogelstein B [6] data on the tissue-specific risk of cancer and lifetime number of stem cell divisions to test whether the number of driver mutations or the mutation rate varies with the lifetime number of stem cell divisions. The driver mutation-initiated carcinogenesis model of Frank SANowak MA [26] is fitted to the data with nonlinear regression. This model incorporates the probability of driver mutations arising during development as well as during adult stem cell division. Using plausible mutation rates, the highest concordance between predicted and observed cancer risk is obtained with single driver mutations across tissues. Testing different fixed numbers of driver mutations, the highest concordance is achieved with single driver mutations and a mutation rate that declines with the lifetime number of stem cell divisions across tissues. Thus, increased inhibition and repair of DNA damage may provide adaptive cancer suppression.

## Methods

### Data

Data on the tissue-specific lifetime risk of cancer (*R*), number of stem cells (*N*), and stem cell division rate (*b*) for 31 tissues are from Tomasetti CVogelstein B [6]. Three cancers whose tissues have *b* = 0 were omitted from analyses. These are glioblastoma, medulloblastoma, and ovarian germ cell cancer. Six cancers were omitted because of conditions that elevate the risk of cancer: colorectal adenocarcinoma with familial adenomatous polyposis (FAP), colorectal adenocarcinoma with Lynch syndrome, duodenum adenocarcinoma with FAP, head & neck squamous cell carcinoma with HPV-16, hepatocellular carcinoma with HCV, and lung adenocarcinoma in smokers.

### Model

The driver mutation-initiated carcinogenesis model of Frank SANowak MA [26] is fitted to the data. In this model, the risk of cancer for a particular stem cell is approximated from the gamma distribution by *R*_*k*_ ≈ (*μ*_*s*_*τ*)^*k*^/*k*!, where *μ*_*s*_ is the driver mutation probability per stem cell division, *τ* is the total number of cell divisions (*τ* = *bT*, where *T* is age), and *k* is the number of driver mutations required to initiate cancer. Driver mutations may also arise during development. The mean frequency of stem cells acquiring a driver mutation during development is 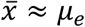 ln(*N*), where *μ*_*e*_ is the driver mutation probability per cell division during exponential growth and *N* is the number of stem cells produced during development to seed the tissue. Therefore, the total risk of cancer is *R*_*T*_ = *N*(1 − *x*)*R*_*k*_ + *NxR*_*k*−1_, where *x* is the frequency of stem cells that start with one driver mutation. This equation was modified to account for the probability that a driver mutation spreads to fixation within a tissue, *ρ*, and the probability that a cancer progresses, *Q* [20]. Thus,

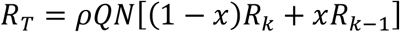

### Nonlinear least squares regression

Numbers of driver mutations, *k*, and stem cell driver mutation rates, *μ*_*s*_, were estimated by fitting the Frank and Nowak model to the data using Levenberg-Marquardt nonlinear least squares regression, as implemented in the R statistical environment [27] package minpack.lm (version 1.2-4) [28]. The quality of the model fit to the data was quantified by the concordance between the predicted and observed cancer risks, calculated using the concordance correlation coefficient, *r*_*c*_, [29] as implemented by the R package DescTools. This is a metric of both correlation and accuracy, while metrics such as root mean squared error (rmse) are strictly for accuracy, and metrics such as *r*^2^ are strictly for correlation. Concordance correlation is superior to coefficients of variation, the paired *t*-test, regression, Pearson correlation, and intraclass correlation for comparing measures [30].

#### Estimating numbers of driver mutations, *k*

To test the hypothesis that the number of driver mutations, *k*, increases with the lifetime number of stem divisions for a tissue, *Nτ*, nonlinear regression was used to estimate values of *k* that explain the lifetime risk of cancer, *R*, across tissues, by fitting the driver mutation-initiated carcinogenesis model of Frank SANowak MA [26] to the data. *k* is assumed to be an allometric (power) function of the lifetime number of stem cell divisions for a tissue: *k* = *a* (*Nτ*)^*c*^, where *a* and *c* are model parameters. Parameter *a* is interpreted as the theoretical value of *k* for a total of one stem cell division (*c* = 0). If *k* increases with *Nτ* at a decreasing rate (hypoallometric), then 0 < *c* < 1. Such a relationship assumes the cost of cancer suppression increases disproportionately with *Nτ*. That is, it should be more costly to suppress cancer as the level of suppression increases, which is reasonable. Estimates for *c* were bounded by −1 and 1.

In estimating numbers of driver mutations, *k* in the Frank and Nowak model was replaced by the right side of the allometric function and then nonlinear regression was used to estimate parameter *c* for the model after fixing parameter *a* and the stem cell driver mutation rate, *μ*_*s*_. This involved finding the value of *a* (at increments of 0.05) that, for a specified value of *μ*_*s*_, gives an estimate of *c* that produces the highest concordance correlation between the predicted and observed lifetime risks of cancer. Both parameters, *a* and *c*, could not be estimated simultaneously.

There are few empirical estimates of the number of driver mutations per stem cell division, *μ*_*s*_. Somatic cell mutation rates have been estimated as ∼10^−9^ point mutations per cell division [31] or ∼10^−7^-10^−6^ per gene per cell division [32-34]. Assuming multiple genes may give rise to driver mutations within a tissue, the mutation rate for driver mutations may be much higher. Bozic I, Antal T, Ohtsuki H, Carter H, Kim D, Chen Set al. [34] estimate a high rate of driver mutations during tumorigenesis of ∼10^−5^ per cell division. Therefore, the range *μ*_*s*_ = 10^−7^-10^−5^ was used in predicting *k*.

Analyses were carried out for each of the three plausible values for *μ*_*s*_ and for probabilities of mutation fixation, *ρ*, and cancer progression, *Q*, (*ρ* = *Q*) of 0.1, 0.01, 0.001, 0.0001, and 0.00001. For simplicity, the mutation rate for exponential growth during development, *μ*_*e*_, was set to equal *μ*_*s*_ (this assumption is relaxed below). For each set of fixed parameters, parameter *c* in the allometric function for *k* was estimated and the function used to calculate minimum and maximum *k* for the minimum and maximum lifetime numbers of stem cell divisions (*Nτ*), respectively, for the tissues analysed.

#### Estimating stem cell mutation rates, *μ*_*s*_

To test the hypothesis that stem cell driver mutation rates, *μ*_*s*_, decrease with the lifetime number of stem divisions for a tissue, *Nτ*, nonlinear regression was used to estimate values of *μ*_*s*_ that explain the lifetime risk of cancer, *R*, across tissues, by fitting the carcinogenesis model of Frank SANowak MA [26] to the data. The model was modified by assuming that *μ*_*s*_ is an allometric function of the lifetime number of stem cell divisions for a tissue: *μ*_*s*_ = *d* (*Nτ*)^*f*^, where *d* is the theoretical mutation rate for a total of one stem cell division (*f* = 0) and −1 < *f* < 0 if *μ*_*s*_ declines with increasing *Nτ* at a decreasing rate, reflecting greater cancer suppression as *Nτ* increases. Estimates for *f* were bounded by –1 and 1.

In estimating stem cell driver mutation rates, *μ*_*s*_ in the Frank and Nowak model was replaced by the right side of the allometric function and then nonlinear regression was used to estimate parameter *f* for the model after fixing parameter *d* and the driver mutation rate for exponential growth during development, *μ*_*e*_, for a given number of driver mutations, *k*. This involved finding the value of *d* (at increments of one order of magnitude) that, for specified values of *k* and *μ*_*e*_, gives an estimate of *f* that produces the highest concordance correlation between the predicted and observed lifetime risks of cancer. Both parameters, *d* and *f*, could not be estimated simultaneously.

When *k* was set to increase with lifetime numbers of stem cell divisions, the value of *k* for each tissue was set by estimating the parameters *a* and *c* for *k* = *a*(*Nτ*)^*c*^ by assigning *k*_min_ to the lowest value of *Nτ* and assigning *k*_max_ to the highest value of *Nτ* and then calculating the slope (*c*) and intercept (log *a*) of the linear relationship on log scales: log *k* = log *a* + *c* log(*Nτ*).

For each assumed number of driver mutations, *k, μ*_*e*_ was set to the three plausible rates (10^−7^, 10^−6^, and 10^−5^). For each set of fixed parameters, parameter *f* in the allometric function for *μ*_*s*_ was estimated and the function used to calculate minimum and maximum *μ*_*s*_ for the maximum and minimum lifetime numbers of stem cell divisions, respectively (mutation rates are expected to decrease: *f* < 0), for the tissues analysed.

## Results

### Numbers of driver mutations, *k*

Nonlinear regression was used to estimate numbers of driver mutations, *k*, that explain the lifetime risk of cancer, *R*, by fitting the carcinogenesis model of Frank SANowak MA [26] to the data and assuming that *k* is an allometric function of the lifetime number of stem cell divisions for a tissue: *k* = *a* (*Nτ*)^*c*^. Highest concordance correlations (*r*_c_ ≥ 0.80) were observed with *ρ* = *Q* = 0.001 and 0.0001 (Table 1). In each case, *k* was estimated to increase with *Nτ* (*c* > 0), but the minimum and maximum values are *k* ≈ 1. Thus, single driver mutations are predicted across tissues. Predictions of *k* > 1 generally have much lower concordance.

**Table 1.** Estimates of the number of driver mutations, with *k* = *a* (*Nτ*)^*c*^ and *μ*_*e*_ = *μ*_*s*_.

| $\mu_s$ | $a$ | Estimate | | $k_{\min}$ | $k_{\max}$ | Concordance | |
| --- | --- | --- | --- | --- | --- | --- | --- |
| | | $c$ | $P$ | | | $r_c$ | 95% CI |
| $\rho = Q = 0.1$ | | | | | | | |
| $10^{-7}$ | 0.85 | 0.0338 | $2.98 \times 10^{-16}$ | 1.39 | 2.25 | 0.60 | 0.29-0.80 |
| $10^{-6}$ | 1.10 | 0.0366 | $3.37 \times 10^{-13}$ | 1.88 | 3.17 | 0.28 | 0.04-0.49 |
| $10^{-5}$ | 2.05 | 0.0297 | $2.55 \times 10^{-11}$ | 3.17 | 4.84 | 0.11 | 0.01-0.21 |
| $\rho = Q = 0.01$ | | | | | | | |
| $10^{-7}$ | 0.70 | 0.0279 | $1.76 \times 10^{-22}$ | 1.05 | 1.57 | 0.79 | 0.56-0.91 |
| $10^{-6}$ | 0.70 | 0.0418 | $2.19 \times 10^{-15}$ | 1.29 | 2.35 | 0.58 | 0.27-0.78 |
| $10^{-5}$ | 1.15 | 0.0233 | $2.42 \times 10^{-12}$ | 1.62 | 2.26 | 0.47 | 0.23-0.66 |
| $\rho = Q = 0.001$ | | | | | | | |
| $10^{-7}$ | 0.55 | 0.0220 | $9.59 \times 10^{-27}$ | 0.76 | 1.04 | 0.81 | 0.60-0.91 |
| $10^{-6}$ | 0.55 | 0.0318 | $1.20 \times 10^{-24}$ | 0.88 | 1.38 | 0.80 | 0.58-0.91 |
| $10^{-5}$ | 0.50 | 0.0565 | $9.00 \times 10^{-14}$ | 1.14 | 2.56 | 0.52 | 0.21-0.74 |
| $\rho = Q = 0.0001$ | | | | | | | |
| $10^{-7}$ | 0.30 | 0.0232 | $1.38 \times 10^{-19}$ | 0.42 | 0.59 | 0.80 | 0.58-0.91 |
| $10^{-6}$ | 0.35 | 0.0266 | $1.09 \times 10^{-22}$ | 0.52 | 0.76 | 0.80 | 0.60-0.91 |
| $10^{-5}$ | 0.35 | 0.0387 | $1.30 \times 10^{-26}$ | 0.62 | 1.07 | 0.81 | 0.60-0.91 |
| $\rho = Q = 0.00001$ | | | | | | | |
| $10^{-7}$ | 0.05 | 0.0355 | $7.64 \times 10^{-07}$ | 0.08 | 0.14 | 0.74 | 0.50-0.88 |
| $10^{-6}$ | 0.05 | 0.0448 | $1.27 \times 10^{-08}$ | 0.10 | 0.18 | 0.75 | 0.51-0.88 |
| $10^{-5}$ | 0.05 | 0.0572 | $6.06 \times 10^{-11}$ | 0.12 | 0.26 | 0.76 | 0.52-0.89 |

### Stem cell driver mutation rates, *μ*_*s*_

Nonlinear regression was used to predict the stem cell driver mutation rates, *μ*_*s*_, that explain the lifetime risk of cancer, *R*, across tissues, by fitting the carcinogenesis model of Frank SANowak MA [26] to the data and assuming that *μ*_*s*_ is an allometric function of the lifetime number of stem cell divisions for a tissue: *μ*_*s*_ = *d* (*Nτ*)^*f*^. Analyses were carried out for *ρ* = *Q* = 0.001 and 0.0001, which give the highest concordance correlations when estimating numbers of driver mutations (Table 1). The highest concordance (*r*_*c*_ = 0.80) is obtained with *k* = 1 for both *ρ* = *Q* = 0.001 and 0.0001 (Table 2; Fig. 1A). With *ρ* = *Q* = 0.001, *μ*_*s*_ is predicted to decline from ∼10^−6^ to ∼10^−8^ with increasing *Nτ* across tissues (*P* < 10^−18^) (Table 2; Fig. 1B). With *ρ* = *Q* = 0.0001, *μ*_*s*_ is predicted to decline from ∼10^−3^ to ∼10^−5^ (*P* < 10^−27^), which is less plausible. Thus, stem cell mutation rates are predicted to decline by two orders of magnitude with increasing lifetime numbers of stem cell divisions. In each case, the minimum stem cell mutation rate 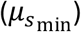 is lower than the corresponding fixed development mutation rate (*μ*_*e*_) (Table 2).

**Table 2.** Estimates of stem cell driver mutation rates, with *μ*_*s*_ = *d* (*Nτ*)^*f*^.

| $k$ | $\mu_e$ | $d$ | Estimate | | $\mu_{s\min}$ | $\mu_{s\max}$ | Concordance | |
| --- | --- | --- | --- | --- | --- | --- | --- | --- |
| | | | $f$ | $P$ | | | $r_c$ | 95% CI |
| $\rho = Q = 0.001$ | | | | | | | | |
| 1 | $10^{-7}$ | $10^{-3}$ | -0.3361 | $3.69 \times 10^{-28}$ | $6.07 \times 10^{-08}$ | $7.33 \times 10^{-06}$ | 0.80 | 0.59-0.90 |
| 1 | $10^{-6}$ | $10^{-3}$ | -0.3648 | $7.19 \times 10^{-19}$ | $2.64 \times 10^{-08}$ | $4.81 \times 10^{-06}$ | 0.80 | 0.61-0.91 |
| 1 | $10^{-5}$ | – | – | – | – | – | – | – |
| 1-2 | $10^{-7}$ | $10^{-4}$ | -0.1164 | $1.42 \times 10^{-08}$ | $3.46 \times 10^{-06}$ | $1.82 \times 10^{-05}$ | 0.68 | 0.38-0.85 |
| 1-2 | $10^{-6}$ | $10^{-5}$ | -0.0310 | $1.33 \times 10^{-02}$ | $4.08 \times 10^{-06}$ | $6.35 \times 10^{-06}$ | 0.69 | 0.41-0.86 |
| 1-2 | $10^{-5}$ | $10^{-5}$ | -0.0306 | $1.13 \times 10^{-02}$ | $4.13 \times 10^{-06}$ | $6.39 \times 10^{-06}$ | 0.74 | 0.48-0.88 |
| 2 | $10^{-7}$ | 1 | -0.4328 | $2.66 \times 10^{-23}$ | $3.70 \times 10^{-06}$ | $1.78 \times 10^{-03}$ | 0.59 | 0.26-0.80 |
| 2 | $10^{-6}$ | 1 | -0.4328 | $2.59 \times 10^{-23}$ | $3.70 \times 10^{-06}$ | $1.78 \times 10^{-03}$ | 0.59 | 0.26-0.80 |
| 2 | $10^{-5}$ | 1 | -0.4296 | $8.53 \times 10^{-25}$ | $4.07 \times 10^{-06}$ | $1.87 \times 10^{-03}$ | 0.63 | 0.31-0.82 |
| $\rho = Q = 0.0001$ | | | | | | | | |
| 1 | $10^{-7}$ | 0.1 | -0.3340 | $9.63 \times 10^{-29}$ | $6.44 \times 10^{-06}$ | $7.56 \times 10^{-04}$ | 0.79 | 0.59-0.90 |
| 1 | $10^{-6}$ | 0.1 | -0.3342 | $1.08 \times 10^{-28}$ | $6.41 \times 10^{-06}$ | $7.54 \times 10^{-04}$ | 0.79 | 0.59-0.90 |
| 1 | $10^{-5}$ | 0.1 | -0.3361 | $3.69 \times 10^{-28}$ | $6.07 \times 10^{-06}$ | $7.33 \times 10^{-04}$ | 0.80 | 0.59-0.90 |
| 1-2 | $10^{-7}$ | 0.1 | -0.2790 | $4.06 \times 10^{-15}$ | $3.16 \times 10^{-05}$ | $1.69 \times 10^{-03}$ | 0.68 | 0.37-0.85 |
| 1-2 | $10^{-6}$ | 0.1 | -0.2790 | $4.03 \times 10^{-15}$ | $3.16 \times 10^{-05}$ | $1.69 \times 10^{-03}$ | 0.68 | 0.37-0.85 |
| 1-2 | $10^{-5}$ | 0.1 | -0.2789 | $3.69 \times 10^{-15}$ | $3.16 \times 10^{-05}$ | $1.69 \times 10^{-03}$ | 0.68 | 0.39-0.85 |
| 2 | $10^{-7}$ | 1 | -0.3570 | $5.90 \times 10^{-18}$ | $3.31 \times 10^{-05}$ | $5.40 \times 10^{-03}$ | 0.52 | 0.24-0.72 |
| 2 | $10^{-6}$ | 1 | -0.3570 | $5.84 \times 10^{-18}$ | $3.31 \times 10^{-05}$ | $5.40 \times 10^{-03}$ | 0.52 | 0.24-0.72 |
| 2 | $10^{-5}$ | 1 | -0.3569 | $5.36 \times 10^{-18}$ | $3.32 \times 10^{-05}$ | $5.40 \times 10^{-03}$ | 0.53 | 0.25-0.73 |

**Figure 1.**
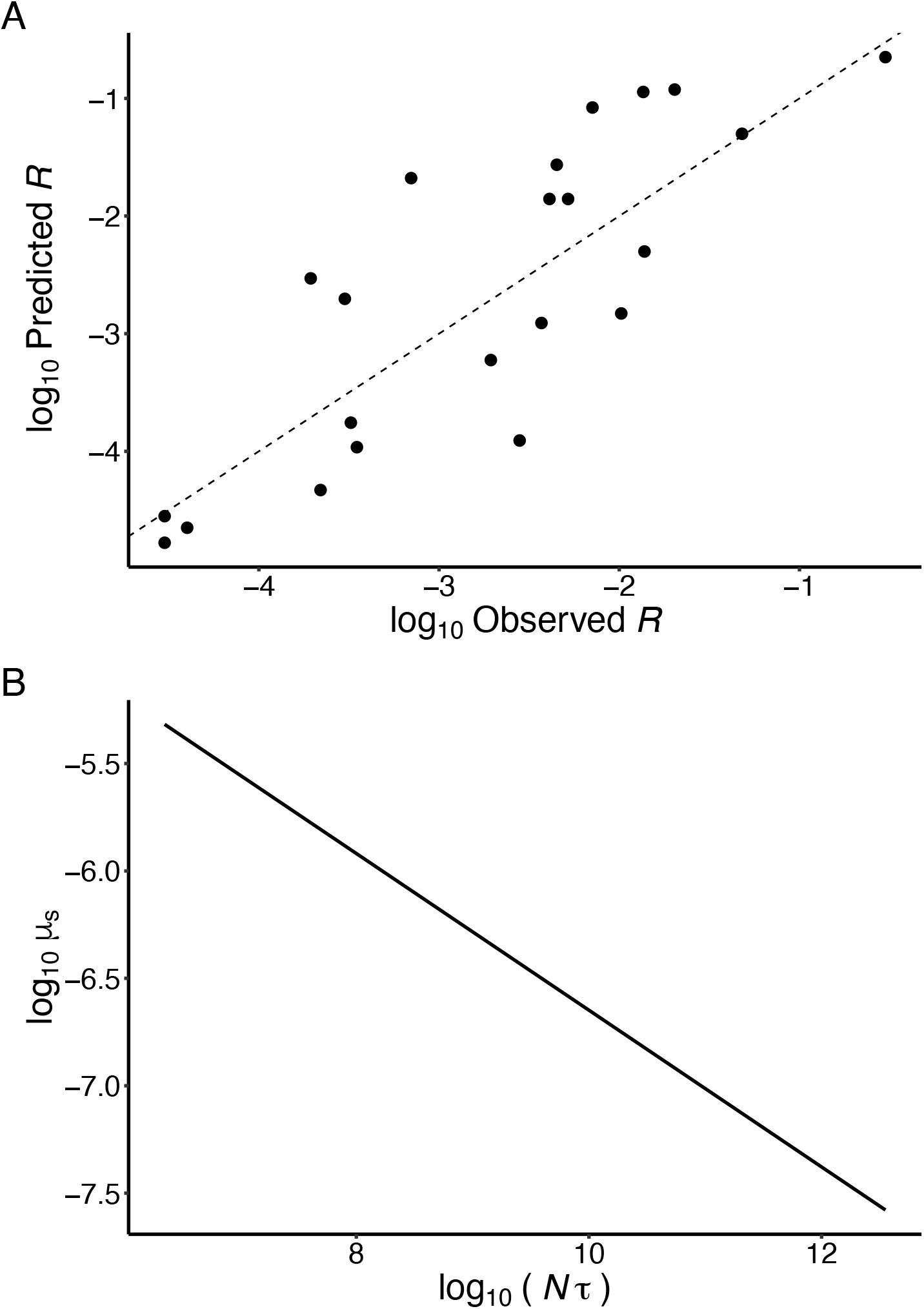
Estimates of stem cell driver mutation rates, *μ*_*s*_. Predicted values are for fixed parameters *ρ* = *Q* = 0.001, *k* = 1, *μ*_*e*_ = 10^−6^, *d* = 10^−3^ and estimated parameter *f* = −0.3648 (*P* = 7.19 × 10^−19^). (A) Predicted and observed lifetime risks of cancer, *R*. Dashed line indicates predicted = observed. Concordance correlation coefficient: *r*_*c*_ = 0.80. (B) Estimated stem cell driver mutation rate as a function of lifetime number of stem cell divisions: *μ*_*s*_ = *d* (*Nτ*)^*f*^.

## Discussion

The results show clear support for single driver mutations across tissues, indicating that the number of driver mutations, *k*, does not increase with the lifetime number of stem cell divisions for a tissue, *Nτ*, as would be expected if *k* is adapted to reduce cancer risk. In contrast, the stem cell driver mutation rate, *μ*_*s*_, is estimated to decrease with increasing *Nτ*. This provide the first evidence of what appears to be differential adaptive cancer suppression within tissues, with stronger suppression in tissues with higher *Nτ*. Stronger cancer suppression may manifest as a lower stem cell mutation rate.

The role of stem cell division in increasing the risk of cancer is supported by the increase in cancer risk with the lifetime number stem cell divisions across tissues [6] and by elevated cancer risk in patients with cancer predisposition syndromes that increase cellular proliferation. An example is familial adenomatous polyposis (FAP), caused by inherited mutations that deactivate the tumour suppressor *APC*. This condition prevents enterocyte precursors from differentiating, retaining stem-like qualities, and thus continuing to divide rather than migrating out of the colonic crypt [35]. The result is a ∼20-fold in increase in the risk of colorectal adenocarcinoma and a ∼100-fold risk of duodenum adenocarcinoma [6]. Infection with some viruses, such as hepatitis C virus (HCV), may also increase cell proliferation associated with chronic inflammation [36]. Hepatocellular carcinoma with HCV has a ∼10-fold increased risk of cancer [6].

Early estimates of numbers of driver mutations necessary to initiate specific cancers ranged from two to eight [3]. A more recent statistical analysis across 30 cancer types infers a mean of only two driver mutations and a range of 1-5 [37]. This estimate closely matches the result of a recent study of 10,478 cancer genomes across 35 cancer types that estimates an overall median of two predicted driver mutations per tumour, with a range of 1-6 [1]. In that study, 51% of tumours had a median of only one driver mutation. These estimates support low numbers of driver mutations, consistent with the estimates in the present study. It should be noted that single driver mutations are consistent with both dominant mutations that activate oncogenes as well as dominant mutations that deactivate tumour suppressor genes with loss of heterozygosity [35].

The fate of new driver mutations is poorly understood. These mutations are generally viewed as being under positive selection [e.g., 38]. Therefore, the fate of a new driver mutation will depend on the strength of selection and genetic drift. Bozic I, Antal T, Ohtsuki H, Carter H, Kim D, Chen Set al. [34] estimated the selection coefficient for driver mutations for glioblastoma multiforme and pancreatic cancers to be *s* = 0.004. This indicates a probability of fixation for a new mutation within the tissue of *ρ* ≈ 2*s* = 0.008. In the present study, high concordance was achieved assuming *ρ* = *Q* = 0.001, which is roughly in line with the estimate of *s*. There do not appear to be any estimates of the probability of progression, *Q*, which may be affected by various factors [34], including tissue architecture [39-41].

Stem cell driver mutation rates, *μ*_*s*_, are predicted to decline from ∼10^−6^ to ∼10^−8^ with increasing *Nτ* across tissues. Thus, maximum *μ*_*s*_ falls within the range of plausible rates (10^−7^-10^−5^). Minimum *μ*_*s*_ is lower than normal rates, which may be explained as an adaptation to high cancer risk. These mutation rates were predicted with developmental mutation rates, *μ*_*e*_, fixed at 10^−7^ and 10^−6^ and thus minimum *μ*_*s*_ is lower than *μ*_*e*_. A recent review suggests that somatic mutation rates may be higher early in development than in adulthood because of rapid cell division or a lack of transcription-coupled DNA repair in early embryos, among other causes [42]. A much higher mutation rate during development compared to adulthood would result in most somatic driver mutations being acquired during development with only a small number acquired during adult stem cell division [26, 43]. The results of the present study, however, support a developmental mutation rate equal to the high adult stem cell mutation rates (*μ*_*e*_ = *μ*_*s*_ = 10^−7^-10^−6^). Thus, low mutation rates adapted to reduce the risk of cancer in tissues with high *Nτ* could explain why mutation rates early in life appear higher than in adulthood.

Somatic mutation rates may be elevated in patients with cancer predisposition syndromes [44], suggesting that mutation rates are selectively maintained at lower levels. For example, the somatic mutation rate in colorectal cancer tumours is ∼8 times higher in patients with an inherited DNA mismatch repair deficiency (Lynch syndrome) than in normal patients, increasing the risk of this cancer ∼10-fold [6]. And, similar rates of somatic mutation accumulation across tissues despite different stem cell division rates [45] suggest that mutation rates per stem cell division are lower in tissues with higher cell division rates.

Somatic mutation rates are under strong selection. This is the implication of the surprisingly little variation in somatic mutation burden and cancer risk among mammal species of vastly different sizes and lifespans [46-49]. For example, although fibroblasts in the bowhead whale, the second largest and longest-lived mammal, require fewer driver mutations for cancer initiation compared to humans, they exhibit enhanced DNA double-strand break repair and fidelity and lower mutation rates compared to fibroblasts of mouse and human [50]. Life history theory explains these observations as the result of investment in costly somatic maintenance, including cancer suppression, that must be optimised to maximise fitness [51]. A similar argument applies across tissues within an individual [21]. Some tissues are more prone to cancer because of large numbers of stem cells or because of high stem cell division rates. Investing more in costly cancer suppression in these tissues than in tissues less prone to cancer could maximize reproductive output by increasing reproductive lifespan while maintaining high levels of investment in reproductive functions. The results of the present study suggest that the mechanism of adaptive cancer suppression across tissues does not involve an increase in the number of driver mutations, but a reduction in the stem cell mutation rate.

## References

1 Kinnersley, B., Sud, A., Everall, A., Cornish, A. J., Chubb, D., Culliford, R., Gruber, A. J., Lärkeryd, A., Mitsopoulos, C., Wedge, D., Houlston, R. 2024 Analysis of 10,478 cancer genomes identifies candidate driver genes and opportunities for precision oncology. Nature Genetics. 56, 1868–1877. (10.1038/s41588-024-01785-9)

2 Jassim, A., Rahrmann, E. P., Simons, B. D., Gilbertson, R. J. 2023 Cancers make their own luck: theories of cancer origins. Nature Reviews Cancer. 23, 710–724. (10.1038/s41568-023-00602-5)

3 Vogelstein, B., Papadopoulos, N., Velculescu, V. E., Zhou, S. B., Diaz, L., Kinzler, K. W. 2013 Cancer Genome Landscapes. Science. 339, 1546–1558. (10.1126/science.1235122)

4 Futreal, P. A., Coin, L., Marshall, M., Down, T., Hubbard, T., Wooster, R., Rahman, N., Stratton, M. R. 2004 A census of human cancer genes. Nature Reviews Cancer. 4, 177–183. (10.1038/nrc1299)

5 Frank, S. A., Nowak, M. A. 2004 Problems of somatic mutation and cancer. Bioessays. 26, 291–299. (10.1002/bies.20000)

6 Tomasetti, C., Vogelstein, B. 2015 Variation in cancer risk among tissues can be explained by the number of stem cell divisions. Science. 347, 78–81. (10.1126/science.1260825)

7 Tomasetti, C., Li, L., Vogelstein, B. 2017 Stem cell divisions, somatic mutations, cancer etiology, and cancer prevention. Science. 355, 1330–1334. (doi:10.1126/science.aaf9011)

8 Nunney, L. 2018 Size matters: height, cell number and a person’s risk of cancer. Proceedings of the Royal Society B: Biological Sciences. 285,

9 Choi, Y. J., Lee, D. H., Han, K.-D., Yoon, H., Shin, C. M., Park, Y. S., Kim, N. 2019 Adult height in relation to risk of cancer in a cohort of 22,809,722 Korean adults. British Journal of Cancer. 120, 668–674. (10.1038/s41416-018-0371-8)

10 Green, J., Cairns, B. J., Casabonne, D., Wright, F. L., Reeves, G., Beral, V. 2011 Height and cancer incidence in the Million Women Study: prospective cohort, and meta-analysis of prospective studies of height and total cancer risk. Lancet Oncology. 12, 785–794.

11 Kabat, G. C., Kim, M. Y., Hollenbeck, A. R., Rohan, T. E. 2014 Attained height, sex, and risk of cancer at different anatomic sites in the NIH-AARP Diet and Health Study. Cancer Causes & Control. 25, 1697–1706. (10.1007/s10552-014-0476-1)

12 Sung, J., Song, Y. M., Lawlor, D. A., Smith, G. D., Ebrahim, S. 2009 Height and Site-specific Cancer Risk: A Cohort Study of a Korean Adult Population. American Journal of Epidemiology. 170, 53–64. (10.1093/aje/kwp088)

13 Wirén, S., Häggström, C., Ulmer, H., Manjer, J., Bjorge, T., Nagel, G., Johansen, D., Hallmans, G., Engeland, A., Concin, H., et al. 2014 Pooled cohort study on height and risk of cancer and cancer death. Cancer Causes & Control. 25, 151–159. (10.1007/s10552-013-0317-7)

14 Kraus, C., Snyder-Mackler, N., Promislow, D. E. L. 2023 How size and genetic diversity shape lifespan across breeds of purebred dogs. GeroScience. 45, 627–643. (10.1007/s11357-022-00653-w)

15 Adams, V. J., Evans, K. M., Sampson, J., Wood, J. L. N. 2010 Methods and mortality results of a health survey of purebred dogs in the UK. Journal of Small Animal Practice. 51, 512–524. (10.1111/j.1748-5827.2010.00974.x)

16 Fleming, J. M., Creevy, K. E., Promislow, D. E. L. 2011 Mortality in North American dogs from 1984 to 2004: An investigation into age-, size-, and breed-related causes of death. Journal of Veterinary Internal Medicine. 25, 187–198. (10.1111/j.1939-1676.2011.0695.x)

17 da Silva, J., Cross, B. J. 2023 Dog life spans and the evolution of aging. The American Naturalist. 201, E000–E000. (10.1086/724384)

18 Doherty, A., Lopes, I., Ford, C. T., Monaco, G., Guest, P., de Magalhães, J. P. 2020 A scan for genes associated with cancer mortality and longevity in pedigree dog breeds. Mammalian Genome. 31, 215–227.

19 Nunney, L. 2024 The effect of body size and inbreeding on cancer mortality in breeds of the domestic dog: a test of the multi-stage model of carcinogenesis. R Soc Open Sci. 11, 231356. (10.1098/rsos.231356)

20 Nowak, M. A., Waclaw, B. 2017 Genes, environment, and “bad luck”. Science. 355, 1266–1267. (10.1126/science.aam9746)

21 Nunney, L., Muir, B. 2015 Peto’s paradox and the hallmarks of cancer: constructing an evolutionary framework for understanding the incidence of cancer. Philosophical Transactions of the Royal Society B: Biological Sciences. 370, 20150161. (doi:10.1098/rstb.2015.0161)

22 Noble, R., Kaltz, O., Hochberg, M. E. 2015 Peto’s paradox and human cancers. Philosophical Transactions of the Royal Society B: Biological Sciences. 370, 20150104. (doi:10.1098/rstb.2015.0104)

23 Noble, R., Kaltz, O., Nunney, L., Hochberg, M. E. 2016 Overestimating the Role of Environment in Cancers. Cancer Prevention Research. 9, 773–776. (10.1158/1940-6207.Capr-16-0126)

24 Tollis, M., Schiffman, J. D., Boddy, A. M. 2017 Evolution of cancer suppression as revealed by mammalian comparative genomics. Current Opinion in Genetics & Development. 42, 40–47. (10.1016/j.gde.2016.12.004)

25 Seluanov, A., Gladyshev, V. N., Vijg, J., Gorbunova, V. 2018 Mechanisms of cancer resistance in long-lived mammals. Nature Reviews Cancer. 18, 433–441. (10.1038/s41568-018-0004-9)

26 Frank, S. A., Nowak, M. A. 2003 Developmental predisposition to cancer. Nature. 422, 494–494. (10.1038/422494a)

27 R Core Team. R: A Language and Environment for Statistical Computing. Vienna, Austria: R Foundation for Statistical Computing 2025.

28 Elzhov, T. V., Mullen, K. M., Spiess, A.-N., Bolker, B. R Interface to the Levenberg-Marquardt Nonlinear Least-Squares Algorithm Found in MINPACK, Plus Support for Bounds. 1.2-4 ed. CRAN 2023.

29 Zar, J. H. 1999 Biostatistical Analysis. 4th ed. Upper Sadle River, NJ: Prentice Hall.

30 Lin, L. I. 1989 A Concordance Correlation-Coefficient to Evaluate Reproducibility. Biometrics. 45, 255-268. (Doi 10.2307/2532051)

31 Lynch, M. 2010 Rate, molecular spectrum, and consequences of human mutation. Proc Natl Acad Sci U S A. 107, 961–968. (10.1073/pnas.0912629107)

32 Hethcote, H. W., Knudson, A. G. 1978 Model for Incidence of Embryonal Cancers -Application to Retinoblastoma. Proceedings of the National Academy of Sciences of the United States of America. 75, 2453–2457. (DOI 10.1073/pnas.75.5.2453)

33 Araten, D. J., Golde, D. W., Zhang, R. H., Thaler, H. T., Gargiulo, L., Notaro, R., Luzzatto, L. 2005 A Quantitative Measurement of the Human Somatic Mutation Rate. Cancer research. 65, 8111–8117. (10.1158/0008-5472.Can-04-1198)

34 Bozic, I., Antal, T., Ohtsuki, H., Carter, H., Kim, D., Chen, S., Karchin, R., Kinzler, K. W., Vogelstein, B., Nowak, M. A. 2010 Accumulation of driver and passenger mutations during tumor progression. Proceedings of the National Academy of Sciences. 107, 18545–18550. (doi:10.1073/pnas.1010978107)

35 Weinberg, R. A. 2014 The Biology of Cancer. Second edition. ed. New York: Garland Science, Taylor C Francis Group.

36 Fiehn, F., Beisel, C., Binder, M. 2024 Hepatitis C virus and hepatocellular carcinoma: carcinogenesis in the era of direct-acting antivirals. Curr Opin Virol. 67, 101423. (10.1016/j.coviro.2024.101423)

37 Iranzo, J., Martincorena, I., Koonin, E. V. 2018 Cancer-mutation network and the number and specificity of driver mutations. Proceedings of the National Academy of Sciences. 115, E6010–E6019.

38 Martincorena, I., Raine, K. M., Gerstung, M., Dawson, K. J., Haase, K., Van Loo, P., Davies, H., Stratton, M. R., Campbell, P. J. 2017 Universal Patterns of Selection in Cancer and Somatic Tissues. Cell. 171, 1029-+. (10.1016/j.cell.2017.09.042)

39 Li, R. Y., Di, L., Li, J., Fan, W. Y., Liu, Y. C., Guo, W. J., Liu, W. L., Liu, L., Li, Q., Chen, L. P., et al. 2021 A body map of somatic mutagenesis in morphologically normal human tissues. Nature. 597, 398-+. (10.1038/s41586-021-03836-1)

40 Lieberman, E., Hauert, C., Nowak, M. A. 2005 Evolutionary dynamics on graphs. Nature. 433, 312–316. (10.1038/nature03204)

41 Nowak, M. A., Michor, F., Iwasa, Y. 2003 The linear process of somatic evolution. Proceedings of the National Academy of Sciences of the United States of America. 100, 14966–14969. (10.1073/pnas.2535419100)

42 Manders, F., van Boxtel, R., Middelkamp, S. 2021 The Dynamics of Somatic Mutagenesis During Life in Humans. Frontiers in Aging. Volume 2 -2021, (10.3389/fragi.2021.802407)

43 Frank, S. A. 2010 Somatic evolutionary genomics: Mutations during development cause highly variable genetic mosaicism with risk of cancer and neurodegeneration. Proceedings of the National Academy of Sciences. 107, 1725–1730. (10.1073/pnas.0909343106)

44 Tomasetti, C., Marchionni, L., Nowak, M. A., Parmigiani, G., Vogelstein, B. 2015 Only three driver gene mutations are required for the development of lung and colorectal cancers. Proceedings of the National Academy of Sciences. 112, 118–123. (doi:10.1073/pnas.1421839112)

45 Blokzijl, F., de Ligt, J., Jager, M., Sasselli, V., Roerink, S., Sasaki, N., Huch, M., Boymans, S., Kuijk, E., Prins, P., et al. 2016 Tissue-specific mutation accumulation in human adult stem cells during life. Nature. 538, 260–264. (10.1038/nature19768)

46 Cagan, A., Baez-Ortega, A., Brzozowska, N., Abascal, F., Coorens, T. H. H., Sanders, M. A., Lawson, A. R. J., Harvey, L. M. R., Bhosle, S., Jones, D., et al. 2022 Somatic mutation rates scale with lifespan across mammals. Nature. (10.1038/s41586-022-04618-z)

47 Peto, R. 2015 Quantitative implications of the approximate irrelevance of mammalian body size and lifespan to lifelong cancer risk. Philosophical Transactions of the Royal Society B: Biological Sciences. 370, 20150198. (doi:10.1098/rstb.2015.0198)

48 Vincze, O., Colchero, F., Lemaitre, J. F., Conde, D. A., Pavard, S., Bieuville, M., Urrutia, A. O., Ujvari, B., Boddy, A. M., Maley, C. C., et al. 2022 Cancer risk across mammals. Nature. 601, 263–267. (10.1038/s41586-021-04224-5)

49 Butler, G., Baker, J., Amend, S. R., Pienta, K. J., Venditti, C. 2025 No evidence for Peto’s paradox in terrestrial vertebrates. Proceedings of the National Academy of Sciences. 122, e2422861122. (doi:10.1073/pnas.2422861122)

50 Firsanov, D., Zacher, M., Tian, X., Sformo, T. L., Zhao, Y., Tombline, G., Lu, J. Y., Zheng, Z., Perelli, L., Gurreri, E., et al. 2025 Evidence for improved DNA repair in the long-lived bowhead whale. Nature. 648, 717–725. (10.1038/s41586-025-09694-5)

51 Caulin, A. F., Maley, C. C. 2011 Peto’s Paradox: evolution’s prescription for cancer prevention. Trends in Ecology & Evolution. 26, 175–182. (10.1016/j.tree.2011.01.002)

